# The algae community in taxon Haptophyceae at the early bloom stage of *Phaeocystis globosa* in Northern Beibu Gulf in winter

**DOI:** 10.1101/492454

**Authors:** Bin Gong, Haiping Wu, Jixian Ma, Meimiao Luo, Xin Li

**Author notes:** Correspondent author: Haiping Wu, Bin Gong.

## Abstract

*Phaeocystis globosa* (Order Phaeocystales, family Phaeocystaceae) caused significant impact on aquaculture farming, global climate change and industry. Since the year 2010, intensive red tides of *P. globosa* began to appear in Beibu Gulf, where previously free of harmful algal blooms, and posed great threats to the cooling systems of a nuclear power plant in 2014 and 2015. In order to discover the bloom mechanism, the community structure of marine microalgae, with a focus on Haptophyceae taxa, in winter in the northern Beibu Gulf near the Qinzhou Bay, Sanniang Bay (SNB) and Dafenjiang River Estuary (DRE), were explored via 18S ribosomal DNA analysis of the V4 region using the Illumina-Based Sequencing platform. The correlation between the relative abundance of five kinds of Haptophyceae algae and environmental factors of seawater were analyzed. The most abundant Haptophyceae-related OTU in terms of number of reads was identified as Phaeocystis and Chrysochromulina. The abundance for other Haptophyceae class was relatively low, such as Haptolina, Prymnesium and Isochrysis. Phaeocystis was present in all samples sites except S6, S11, S12, S14 and S15, and particularly abundant at S8, nearly 29 times more than the second most abundant site. Most notably, the results showed that Phaeocystis displayed highly positive linear correlation with the concentration of NO_3_^-^-N (Pearson r=0.856, p<0.01). Linear regression analysis indicated that Phaeocystis was significantly linearly related to the NO_3_^-^-N (R2=0.732; Y=-0.005 + 0.410*X, Y is the relative abundance of *P.globosa*, X is the concentration of NO_3_^-^-N; F=38.227, P<0.05) and NO_3_^-^-N has a significant positive effect on *P.globosa* (regression coefficient is 0.410, P=0.000). Moreover, the relative abundance of Phaeocystis was significant related to temperature of sea water (Pearson r=-0.882, p<0.01). Water temperature can explain the 77.8% change reason for the *P.globosa* (R^2^=0.778), and has a significant effect on *P. globosa* (Y=0.169-0.009*X, F=49.031, P<0.05), and the regression coefficient is -0.009 (P=0.000) which indicated a significant negative impact relationship between them. Our high throughput sequencing (HTS) based research illustrated how the *P. globosa* bloom generated and its relationship with NO_3_^-^-N and temperature of sea water in northern Beibu Gulf for the first time, and bringing hope for solving this big problem.

## Introduction

Haptophyceae (or Prymnesiophyceae), a class of the phylum Chrysophyta, contained the Order Prymnesiales, Discoasterales, Phaeocystales, Isochrysidales. The bloom of some Haptophyceae algae had occurred frequently worldwide and led to great ecological disaster and substantial economic losses. For instance, *Prymnesium parvum*, a species of Haptophyceae algae (Order Prymnesiales, family Prymnesiaceae), is capable of producing a toxin, prymnesin, and kills fish [1]. Similarly, large area bloom of the *Chrysochromulina polylepis* (Order Prymnesiales, family Prymnesiaceae) have resulted in mortality of trout and salmon in Scandinavian waters during Spring 1988 [2]. Further more, the bloom of *Phaeocystis globosa* (Order Phaeocystales, family Phaeocystaceae) caused mortality of cultured fish [3], Mussel Mortalities [4], higher concentration of DMS [5] and Clogging of Cooling System of Power Plant [6], so both significantly impact aquaculture farming, global climate change and industry.

Beibu Gulf is an important habitat for protection and propagation of marine organisms, known for Indo-Pacific humpback dolphins [7], horseshoe crabs [8], and also an ecologically sensitive region [9]. Recently, rapid economic development and human activities had already resulted in great degradation of marine environment [10,11]. Most noticeably, human-induced nutrient enrichment is becoming a serious problem for coastal marine areas of Beibu Gulf [9,12]. Since the year 2010, intensive red tides of *P. globosa* began to appear in Beibu Gulf, where previously free of harmful algal blooms, and posed great threats to the cooling systems of a nuclear power plant in 2014 and 2015 [6]. Up to now, knowledge on where the *P. globosa* originated and the bloom mechanism is still quite limited. In our opinion, in order to solve the great confusion above, special attention should be paid to the early bloom stage and the environmental characteristics leading to the bloom. For this purpose, many methods had been utilized, such as automatic monitoring buoy [13], satellite remote sensing [14], “molecular probes” [15], methods to detect and quantify toxins [16] and a combination of methods above [17].

Environmental DNA-based Techniques (EDT) was utilized in monitoring Prokaryote and Eukaryotes in water environments for many years and significantly gained impetus over traditional approaches presently [18]. Recently, there are many EDT methods advented for this object; for instance, DNA metabarcoding [19,20], Microsatellite DNA marker [21], and high throughput sequencing (HTS) techniques based metagenomics [22,23] and amplicons [24,25]. In recent years, the HTS techniques has been widely used and remarkably promoted the ecological studies of bacteria [26], fungi [27], algae [28] and animals [29]. However, to date, molecular diversity of microeukaryotes, including algae, in the Beibu Gulf marine region, remains unexplored. Thus, the recent study about the community and diversity structure of marine microalgae, with a focus on Haptophyceae taxa in winter in the northern Beibu Gulf near the Qinzhou Bay, Sanniang Bay (SNB) and Dafenjiang River Estuary (DRE), were explored via 18S ribosomal DNA analysis of the V4 region using the Illumina-Based Sequencing platform. This study provides a valid taxonomic reference dataset for future microeukaryotic community structure and diversity studies, aimed at monitoring environmental change in the northern Beibu Gulf.

## Materials and Methods

### Sample Collection

The samples were collected on 27 Dec 2017, when large Phaeocystis bloom was observed between eastern Qinzhou Bay (EQB) and Dafenjiang River Estuary (DRE), during the middle several days of Jan to the end of Mar in 2018. Seawater was collected from surface, middle and deep using a CTD Rosette Water Sampling System (Sea-Bird Electronics, USA) between eastern Qinzhou Bay (EQB) and Dafenjiang River Estuary (DRE) (21°32’58’’N, 108°39’56’’E-21°37’39’’N, 108°55’57’’E) and mixed evenly. A total of 16 seawater samples (S1-S16, Figure 1) were collected in 3L sterile polyethylene bottles, kept in the dark at 4-8 ˚C, and filtered at laboratory within 4h. Each 3000 ml seawater sample was filtered through a 1.2 µm mixed cellulose membrane filter (Advantec, Japan) and filtrates were frozen immediately at -20˚C for subsequent molecular analyses.

**Figure 1.**
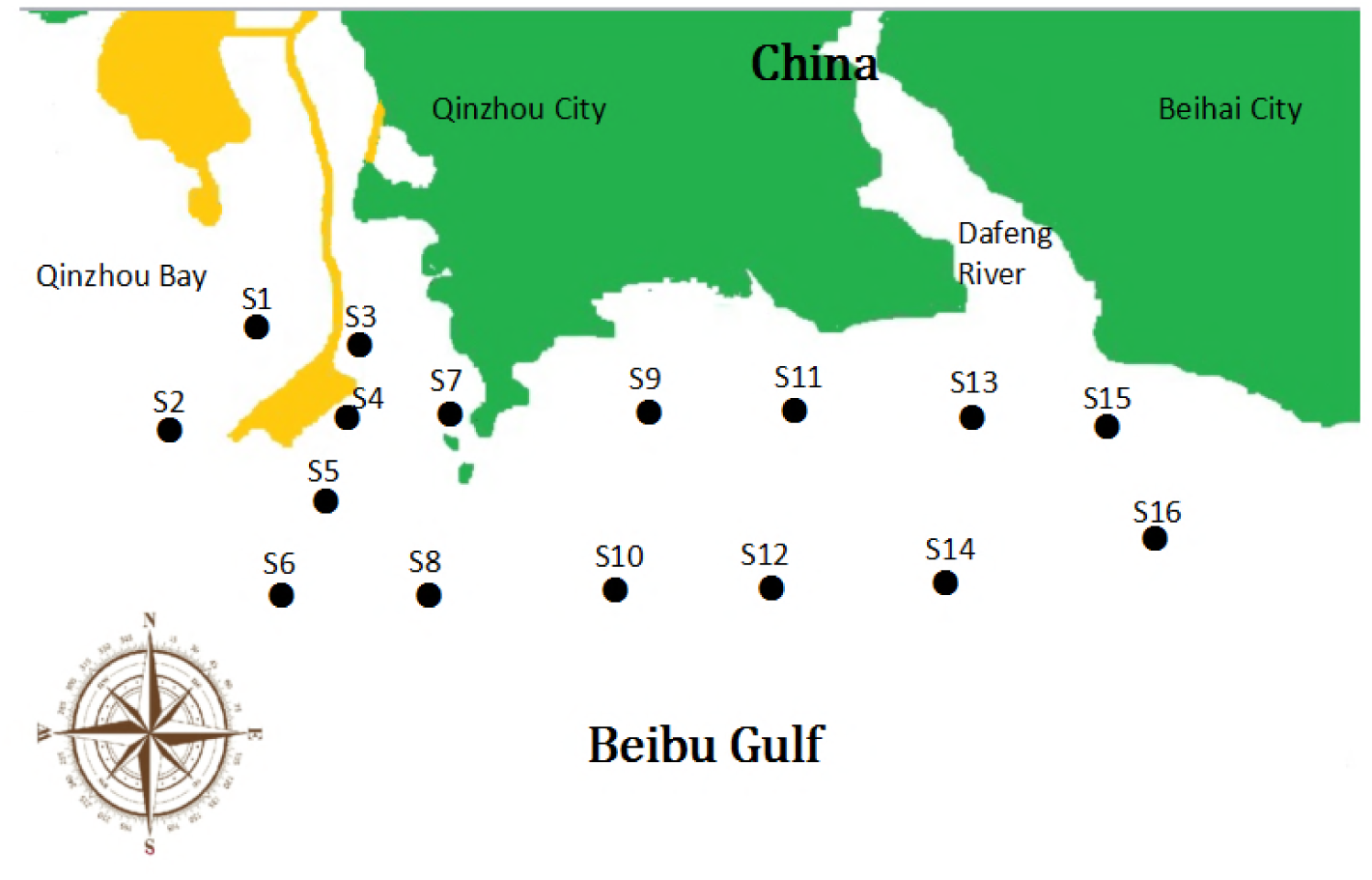
Locations of the the 16 sites (S1-S16) where seawater samples were collected on 27 Dec 2017, including six sites near Sandun Dock (S1-S6), six sites near Sanniang Bay (SNB, S7-S12) and four sites in the Dafengjiang River Estuary (DRE,S13-S16).

### DNA Extraction, PCR and illumina sequencing

The PowerWater DNA isolation kit (MoBio Laboratories Inc., CA, USA) was used to extract the DNA of the total organisms on the 1.2 µm filters following the manufacturer’s protocol. The DNA samples were detected by 1% agarose gels and NanoDrop One spectrophotometer (Thermo Fischer Scientific Inc., USA), and then were amplified using the primers 528F and 706R [30,31] which was designed to amplify the hypervariable region V4 of eukaryote 18S rRNA gene. Illumina sequencing was carried out by the Novogene Company (Beijing, China). Sequencing libraries were generated using TruSeq^®^ DNA PCR-Free Sample Preparation Kit (Illumina, USA) following manufacturer’s recommendations and index codes were added. The library quality was assessed on the Qubit@ 2.0 Fluorometer (Thermo Scientific) and Agilent Bioanalyzer 2100 system. At last, the library was sequenced on an Illumina HiSeq2500 platform and 250 bp paired-end reads were generated. The raw data sequences were assigned to samples by their unique barcodes. The 18S rDNA primers and barcodes were cut off to generate pair-end (PE) reads. Paired-end reads were merged using FLASH (V1.2.7) [32]; the raw tags were filtered to obtain the high-quality clean tags using QIIME software package (V1.7.0) [33]. Sequences analysis were performed by Uparse software (Uparse v7.0.1001) [34]. Sequences with *≥*97% similarity were assigned to the same OTUs. For each representative sequence, the GreenGene Database [35] was used to annotate taxonomic information.

### Physical and chemical analyses of seawater characteristics

The seawater temperature (˚C), salinity were measured using the SBE 911 plus CTD (Sea-Bird Electronics, USA). Dissolved oxygen (DO) concentrations (ml/l) were measured using the SeaBird 43 (Sea-Bird Electronics, USA). The pH values were measured by pH meter (METTLER,FE38-Meter,30254110). Chlorophyll-a fluorescence was measured using the WETStar (WET Labs, USA). Inorganic nutrient concentrations (nitrate [NO_3_^-^-N], ammonia [NH_4_^+^-N], nitrite [NO_2_^-^], and phosphate [PO_4_^3-^]) were determined from 100 mL samples with an Alliance Integral Futura Autoanalyzer II [36,37]. The TOC content was determined with TOC analyzer (Multi N/C 3100, Analytik Jena AG, Jena, Germany) according to the procedure explained by Ali [38]. Spatial distribution of seawater characteristics was interpolated by the measurements of the 16 sampling sites using Kriging method [39]. Interpolations outside the sampling area and over the terrestrial landscape were subtracted.

### Statistical analysis

Data were compiled and transformed in Microsoft Excel. Correlation between variables were made using a linear Pearson’s r coefficient. Linear regression analysis was facilitated and conducted between closely related. Statistics were generated using the SigmaStat version 2.01 software package (SPSS, Inc., Chicago, Ill). All comparisons were performed at the 95% confidence level.

## Result

### OTUs profile based on 18s RNA amplicon analysis using high throughput sequencing (HTS)

The eukaryote communities in the seawater were represented by 1,594 OTUs identified in our present study. A good coverage of the eukaryotic diversity in all samples were illustrated by rarefaction curves, which were calculated and reached a plateau in all cases. All OTUs were classified into 44 main Phylum, and the most abundant were Arthropoda, Ascomycota, Chlorophyta, Diatomea, Basidiomycota, Protalveolata and Picozoa (Fig 2). Among all the OTUs, our most concerned OTU was that identified as class Haptophyceae, which some Harmful algae (such as *P.globosa* and Prymnesium) causing serious ecological disasters represented in this group. In our study, 19 OTUs were classified into Haptophyceae, belonging to the groups of Prymnesium, Haptolina, Isochrysis, Chrysochromulina and Phaeocystis respectively (Fig 3). Phaeocystis was represented by OTU 174 and OTU 1030, with OTU 174 identified as *P. globosa* (100% similarity). Prymnesium was represented only by one OTU (OTU 1269), which closely related to *Prymnesium zebrinum* (98% similarity) and *Prymnesium pienaarii* (97% similarity). The most biggest taxonomic group in Haptophyceae was identified as genus Chrysochromulina which including 4 OTUs. The other taxonomic groups belonging to Haptophyceae were *Haptolina fragaria* (100% similarity, OTU370), *Isochrysis galbana* (100% similarity, OTU1449), *Haptophyceae* sp. (96% similarity, OTU1155) (Fig 3).

**Figure 2.**
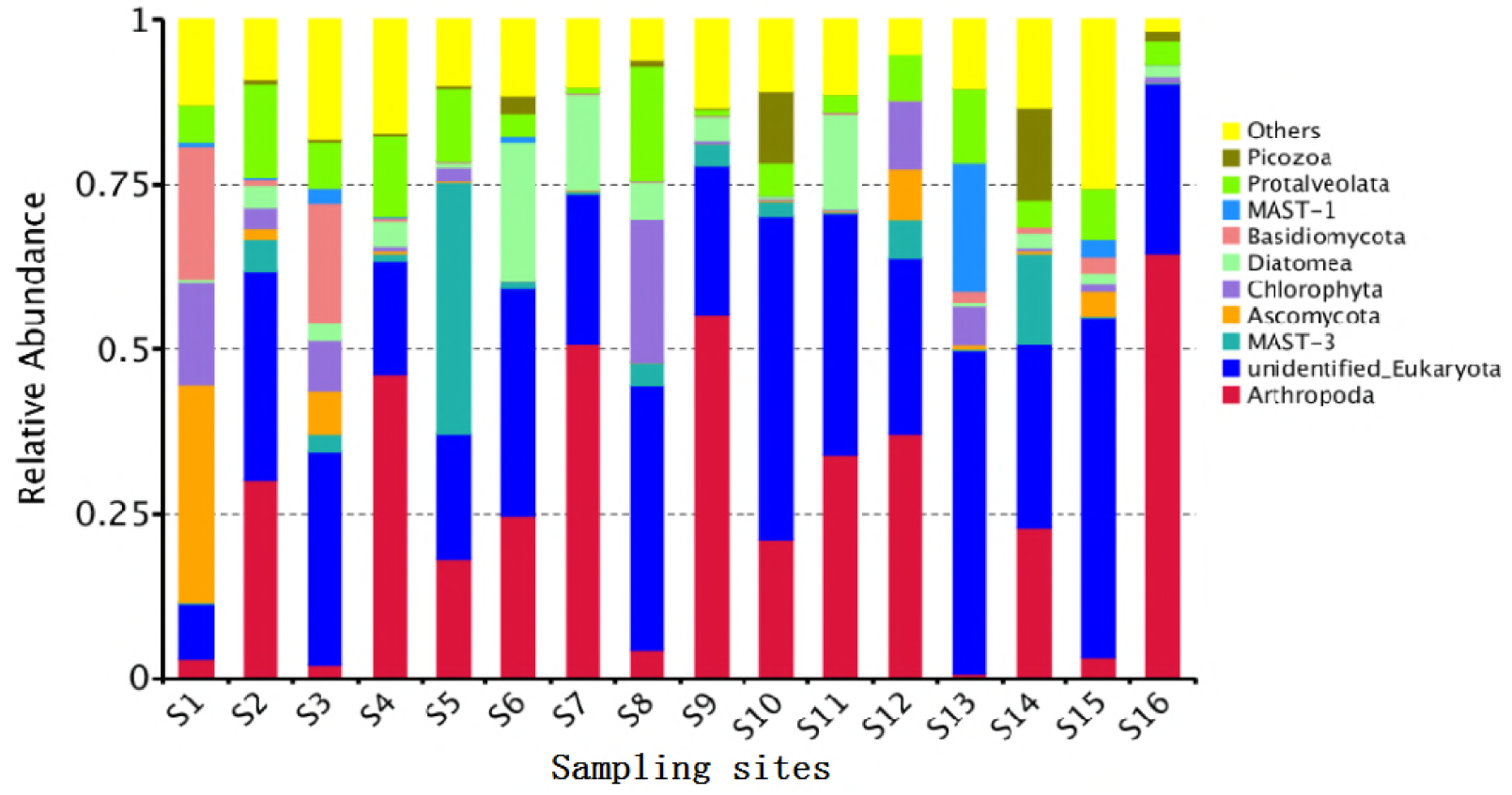
The top ten OTUs were identified based on the hypervariable region V4 of eukaryote 18S rRNA gene and sequences with *≥*97% similarity were assigned to the same OTUs.

**Figure 3.**
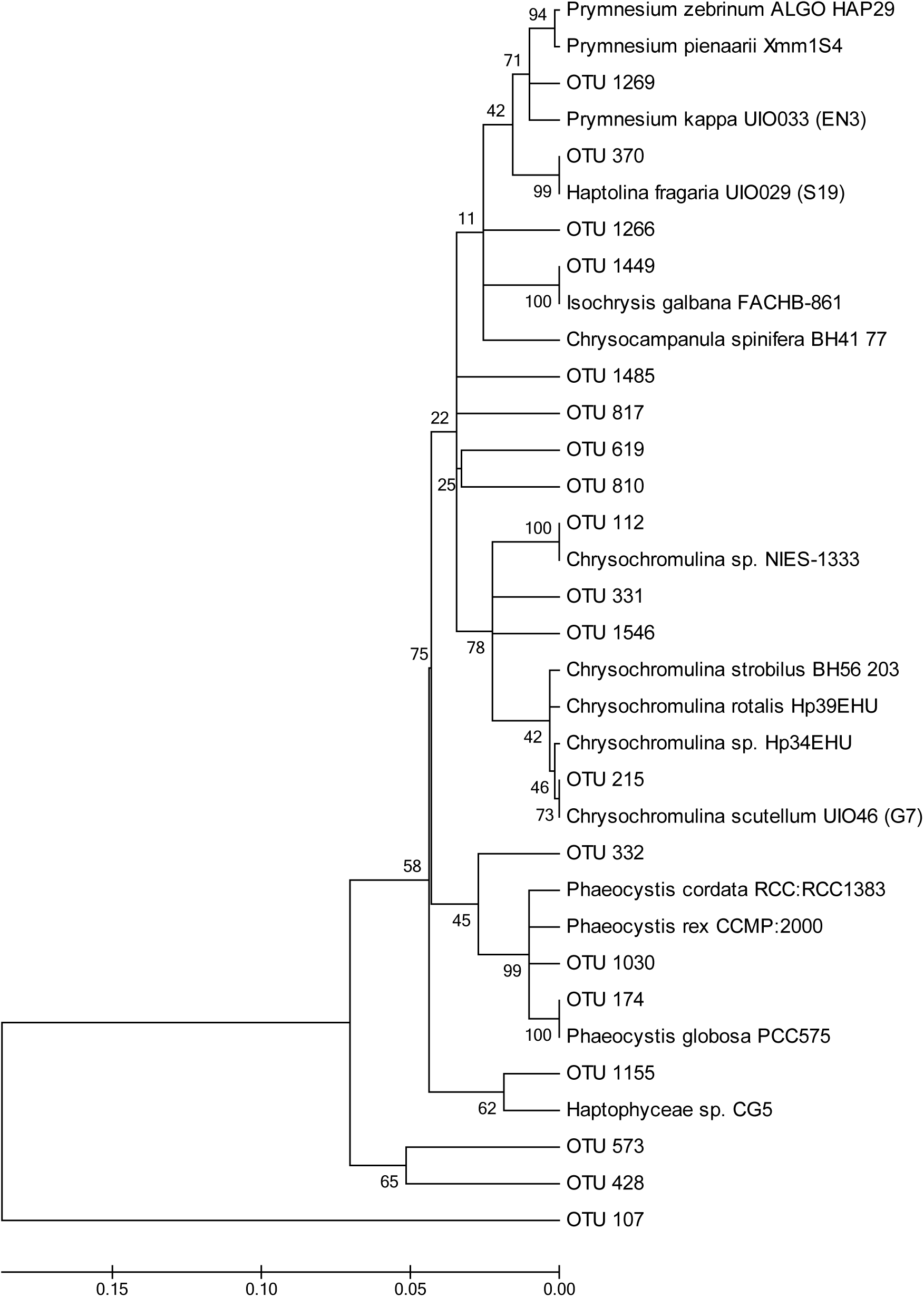
The taxonomic relationship of 18S rDNA phylotypes of class Haptophyceae from the the northern coast of Beibu Gulf. Kimura two-parameter model and midpoint rooting options were used to reconstruct phylogenetic relationships. Numbers above branches indicate bootstraps for NJ analysis (> 50).

The most abundant Haptophyceae-related OTU in terms of number of reads was identified as Phaeocystis and Chrysochromulina. Chrysochromulina was the most abundant taxa containing 5353 reads, while Phaeocystis displayed a relatively lower number of 2298 reads. The abundance for other Haptophyceae class was relatively low, such as Haptolina (228 reads), Prymnesium (9 reads), Isochrysis (6 reads) (Fig 4). The relative abundance of the five kinds of Haptophyceae at different sampling sites showed different characteristics. The spatial distribution of Phaeocystis and Prymnesium were similar. Phaeocystis was present in all samples sites except S6, S11, S12, S14 and S15, and particularly abundant at S8 (2150 reads, 3.4%), nearly 29 times more than the second most abundant site S2 (73 reads). Prymnesium showed the same pattern, recording the highest percentage at site S8 (7 reads, 0.01%) and haven’t appeared on the other sites. While Isochrysis indicated very different property, with major peaks occurred at the sites S3 (5 reads) and none at the other sites. Most of algal species Chrysochromulina were abundant and fairly distributed at different stations with reads ranged from 158 to 899, except highest percentage (2.05 %, Site 8) and relatively lower at S6 (0%), S7 (19 reads, 0.03%), S9 (46 reads, 0.07%), S11 (30 reads, 0.04%), S12 (0%) and S16 (54 reads, 0.08%). The relative abundance of Haptolina among different sites displayed similar pattern with Chrysochromulina, and the major peaks occurred at sites S8 (Fig 5).

**Figure 4.**
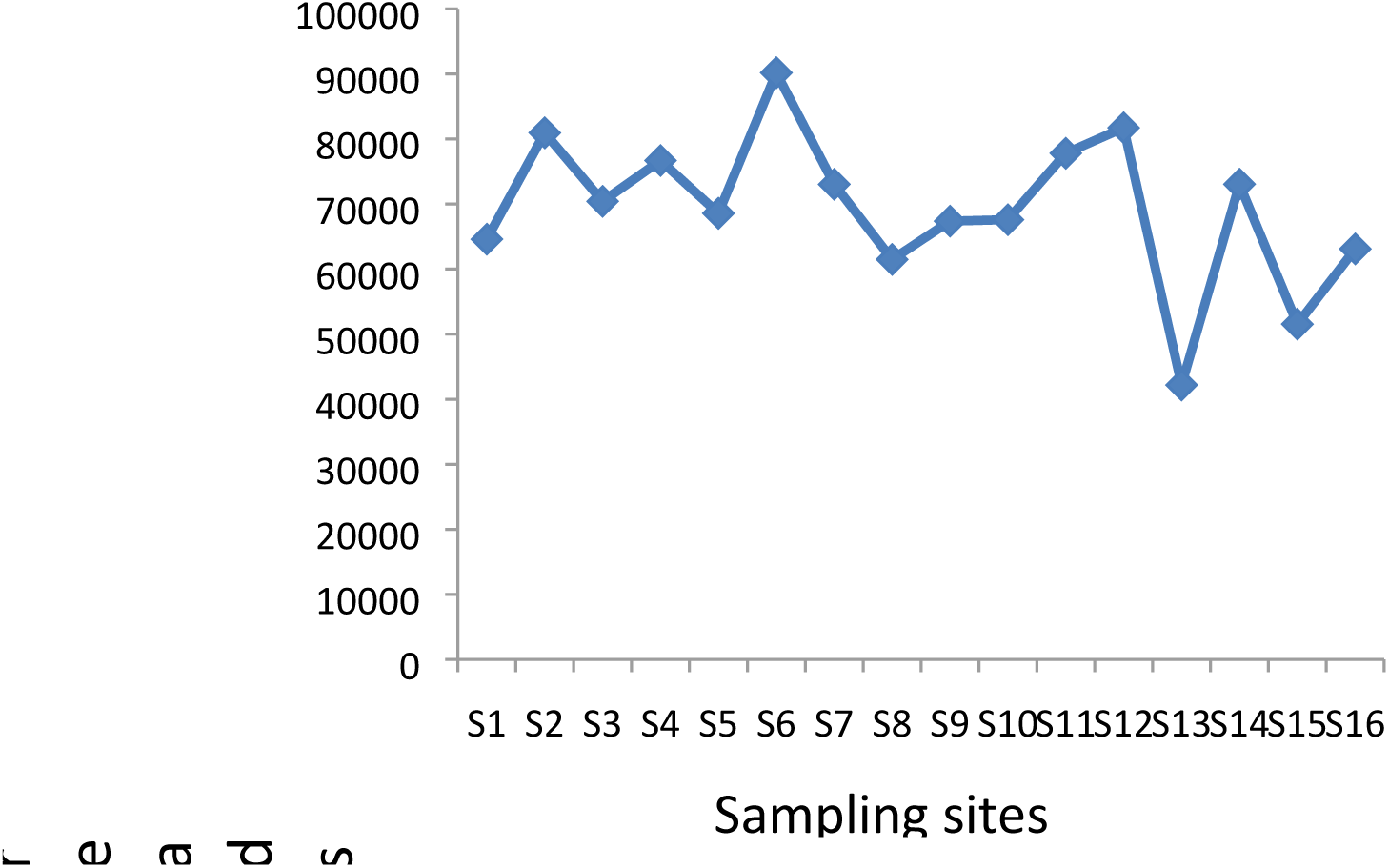
The number of reads detected in 16 sampling sites using 18s RNA amplicon analysis.

**Figure 5.**
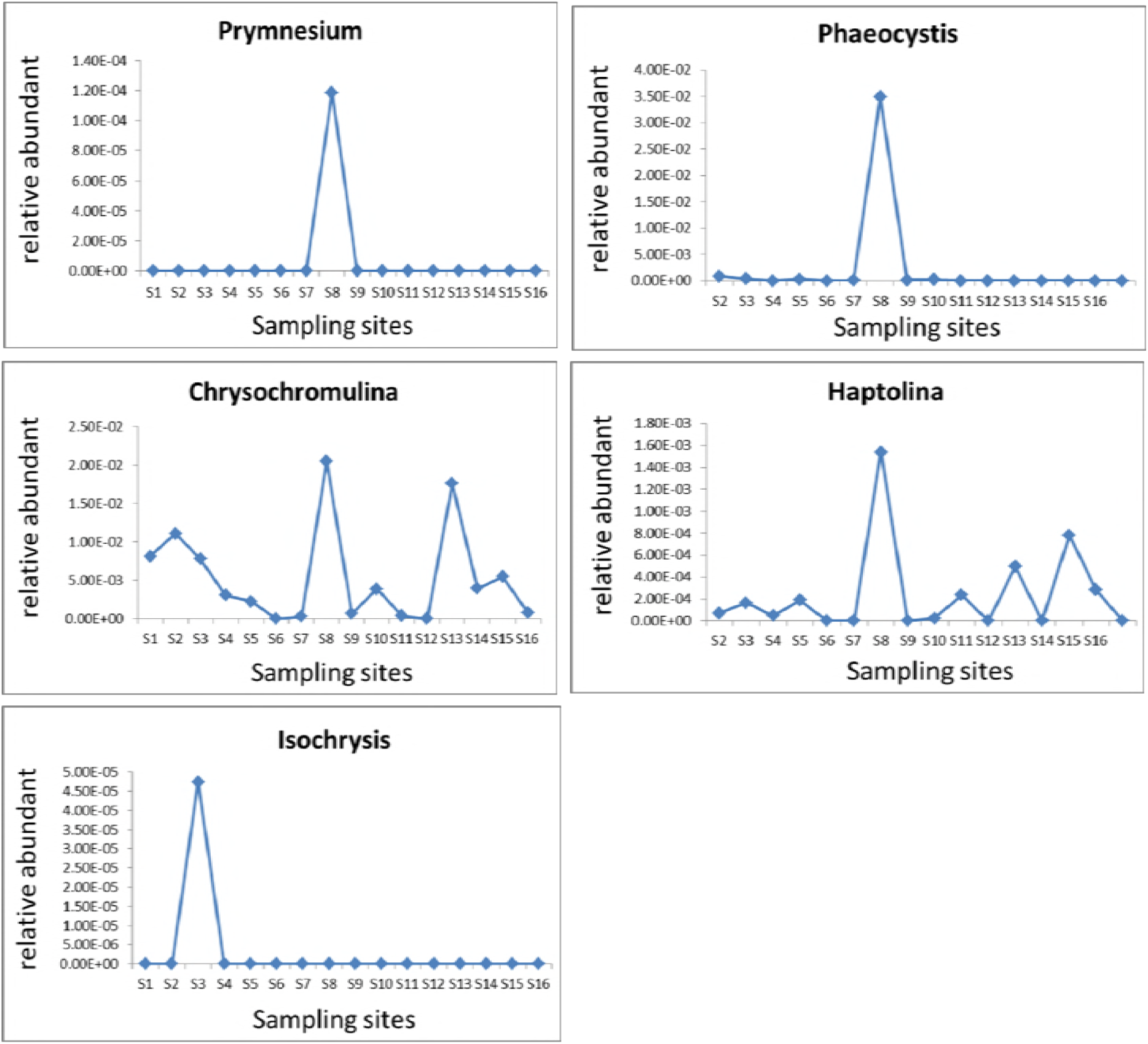
Spacial variations of relative abundance (%) of five Haptophyceae algae (Prymnesium, Haptolina, Isochrysis, Chrysochromulina and Phaeocystis) at the different stations in the northern coast of Beibu Gulf in Dec 2017.

### Physical and chemical analyses of seawater characteristics

Physical and chemical characteristics of the seawater in the 16 sampling sites, including nutrient concentrations (TOC, NH_4_^+^-N, NO_3_^-^-N, NO_2_^-^-N, and phosphate [PO_4_^3-^]) and orther environmental condition (chlorophyll-a, DO, salinity, pH, temperature of seawater), were detected. Just as our above discovery that relative abundance of some Haptophyceae algae were obviously higher at site S8 than others, some environmental factors at S8 were apparently special as well. Seawater temperature during the period of the study ranged from 15.4 to 18.5 ˚C and the lowest temperature appeared at site S8, while the salinity ranged from 25.7 to 37.3 ppt and S8 contained the lowest value 25.69 ppt. The highest value recorded for DO was 9.32 mg/L at site S8 and lowest was 8.33 mg/L at site S13. The highest value of pH was 8.71 at site 12 and lowest was 8.22 at S8. For the nutrient concentrations, the site S8 possessed the highest concentration of NO_3_^-^-N (0.066 mg/L). Spatial distribution of seawater characteristics over the study region, including DO, NO_3_^-^-N, temperature of seawater and pH, were simply illustrated on the map by using blue and red colour (Figure 6).

**Figure 6.**
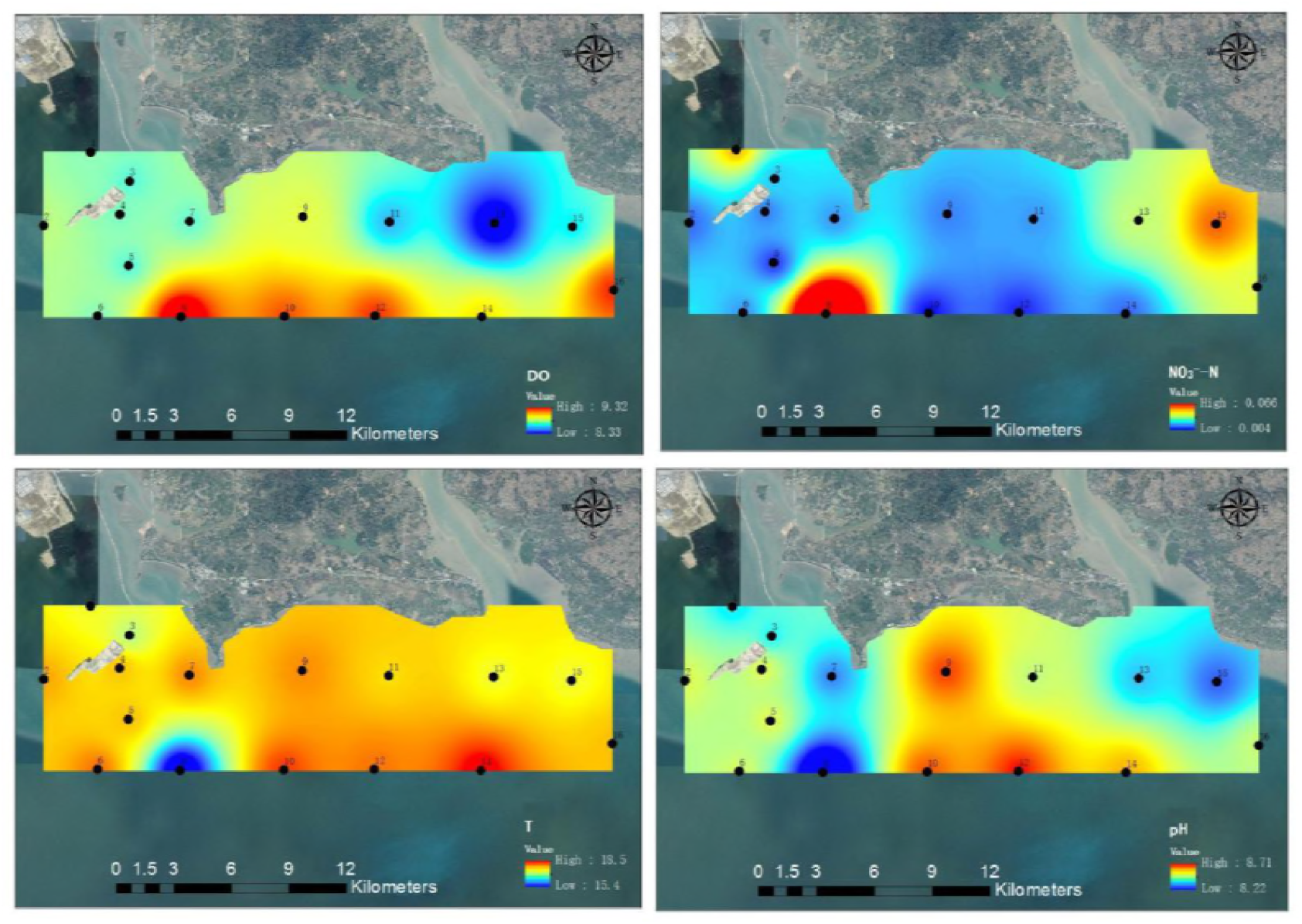
Spatial distribution of seawater characteristics over the study region, including DO (a), NO_3_^-^-N (b), T-temperature of seawater (c) and pH (d).

### The Correlation between environmental factor and Haptophyceae algae

The correlation between the relative abundance of five kinds of Haptophyceae algae and environmental factors of seawater (temperature, pH, salinity, DO, NO_3_^-^-N, NO_2_^-^-N, NH_4_^+^-N, Chlorophyll a and TOC) were analyzed. The results showed that Prymnesium, Phaeocystis and Haptolina displayed highly positive linear correlation with the concentration of NO_3_^-^-N (Pearson r=0.85˜0.92, p<0.01). Isochrysis and Chrysochromulina indicated high negative correlation with temperature of sea water (Pearson r=-0.88˜-0.89, p<0.01). Except for this, there was a significant negative correlation just for Prymnesium, Haptolina, Chrysochromulina and Phaeocystis with pH and salinity (Pearson r=-0.52˜-0.78, p<0.01). Chrysochromulina revealed significant negative correlation with seawater temperature (Pearson r=-0.787, p<0.01). Haptolina and Chrysochromulina also exhibited significant negative correlation with Chlorophyll-a (Pearson r=-0.52∼-0.53, p<0.01). There was a significant positive correlation of Prymnesium and Phaeocystis with DO (Pearson r=0.534, p<0.01), and also the same relationship of Chrysochromulina with NO3^-^-N (Pearson r=0.69, p<0.01). There was no other obvious correlation between other environmental factors and those five kinds of Haptophyceae algae (Fig 7).

**Figure 7.**
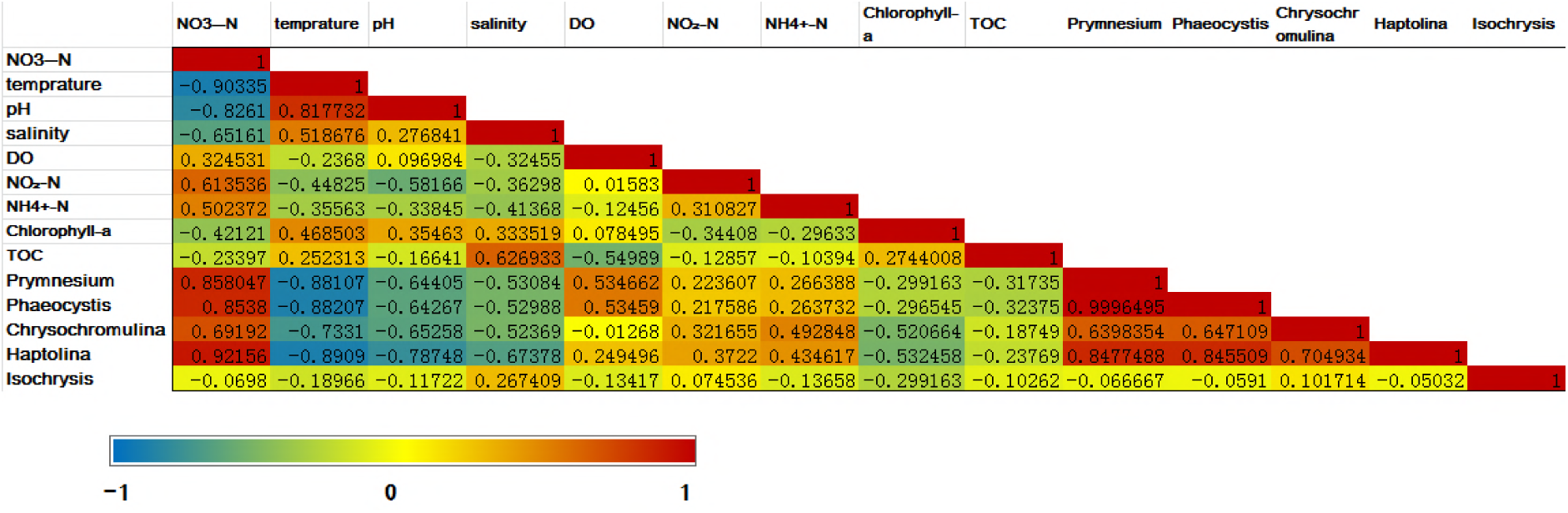
The correlation between the relative abundance of five kinds of Haptophyceae algae and environmental factor (NO_3_^-^-N, temprature, H, salinity, DO, NO2^-^-N, NH4-N, Chlorophyll a and TOC) was indicated by Correlation matrix (Pearson’s product moment correlation coefficient). The more the colure of shaded cells was close to red, the more significantly the correlation was positively related (P < 0.05); inversely, the more the colure of shaded cells was close to blue, the more significantly the correlation was negatively related (P < 0.05).

Chrysochromulina was the most abundant taxon of Haptophyceae algae in this study. The environmental factors NO_3_^-^-N, pH and seawater temperature on the relative abundance of Chrysochromulina were studied by regression analysis. The concentration of NO_3_^-^-N possessed significant positive impact on the relative abundance of Chrysochromulina (Pearson r=0.692, p<0.01; R^2^=0.479; Y=-0.001 + 0.281*X, Y is the relative abundance of Chrysochromulina, X is the concentration of NO3^-^-N; F=12.859, P<0.05; regression coefficient is 0.281, P=0.003). Conversely, the temperature of seawater indicated obvious negative influence on the relative abundance of Chrysochromulina (Pearson r=-0.733, p<0.01; R^2^=0.537; Y=0.123-0.007*X, Y is the relative abundance of Chrysochromulina, X is the value of seawater temperature; F=16.266, P<0.05; regression coefficient is -0.007, P=0.001), and pH showed the same features on it (Pearson r=-0.653, p<0.01; R^2^=0.426; Y=0.282-0.032*X, Y is the relative abundance of Chrysochromulina, X is the value of seawater pH; F=10.384, P<0.05; regression coefficient is -0.032, P=0.006).

There is consistent relation between the relative abundance of Phaeocystis and environmental factor NO_3_^-^-N, DO, pH and temperature of sea water. The relative abundance of Phaeocystis has obvious correlation with NO3^-^-N (Pearson r=0.856, p<0.01). Linear regression analysis indicated that Phaeocystis was significantly linearly related to the NO3^-^-N (R^2^=0.732; Y=-0.005 + 0.410*X, Y is the relative abundance of P.globosa,X is the concentration of NO3^-^-N; F=38.227, P<0.05) and NO3^-^-N has a significant positive effect on *P.globosa* (regression coefficient is 0.410, P=0.000). The relative abundance of Phaeocystis had a positive correlation with DO (Pearson r=0.535, p<0.01; R2=0.286; Y=-0.140 + 0.016*DO,F=5.616, P<0.05; regression coefficient is 0.016,P<0.05). The relative abundance of Phaeocystis was significant related to temperature of sea water (Pearson r=-0.882,p<0.01). Water temperature can explain the 77.8% change reason for the P.globosa (R^2^=0.778), and has a significant effect on P. globosa (Y=0.169-0.009*X, F=49.031, P<0.05), and the regression coefficient is -0.009 (P=0.000) which indicated a significant negative impact relationship between them. The relative abundance of Phaeocystis also has negative correlation with pH of sea water (Pearson r=-0.643, p<0.01).

Prymnesium exhibited very similar condition with Phaeocystis, concerning relationship between their relative abundance and environmental characteristics. For example, the relative abundance of Prymnesium showed highly correlation with NO3^-^-N (Pearson r=0.858, p<0.01) and NO3^-^-N had a significant positive impact on the relative abundance of Prymnesium (R^2^=0.736; Y=-0.000 + 0.002*X, Y is the relative abundance of Prymnesium, X is the concentration of NO3^-^-N; F=39.079, P<0.05; regression coefficient is 0.002, P=0.000). In contrast, the temprature of seawater exhibited obvious negative effect on Prymnesium (R^2^=0.776; Y=0.001 - 0.000*X, Y is the relative abundance of Prymnesium,X is the value of temperature of seawater; F=48.579, P<0.05; regression coefficient is- 0.000, P=0.000).

## Discussion

### NGS: a promising approach to study the community of algae in marine environments

*P.globosa* bloom occurred in winter from the year of 2014-2015 near the coast of Beibu Gulf annually. In ordor to discover the bloom mechanism, *P. globosa* was monitored between eastern Qinzhou Bay (EQB) and Dafenjiang River Estuary (DRE) during Nov 15^th^ 2017 to Feb 15^th^ 2018. The beginning of the *P. globosa* bloom appeared from the middle several days of Jan in 2018, and disappearing occurred at the end of Mar in 2018. All the above findings were based on microscopy observations and cell counts, and it seems that the bloom appeared suddenly and instantly. We believed that finding out what happened before the emergence of *P. globosa,* not only by observations of microscopy and naked eyes, was very significant for our understanding of bloom mechanism of *P. globosa* in Beibu Gulf.

Classically, the algae ecological value is weighted based on the relative abundance of morphologically identified species. This traditional method is costly, time-consuming, and requires excellent taxonomic expertise, which is not always available [40]. Comparatively speaking, the eDNA and NGS approach for identification and quantification of algae open a new avenues for assessing and monitoring of aquatic ecosystems [41]. In our present work, by utilizing NGS based eDNA detection, not only the *P. globosa*, but also all the algae species belonging to Haptophyceae, taxonomically including *P. globosa*, were discovered and analyzed. The two most abundant OTUs in taxon Haptophyceae were affiliated with Chrysochromulina (5353 reads) and Phaeocystis (2298 reads), significantly more reads than Haptolina (228 reads), Prymnesium (9 reads) and Isochrysis (6 reads). The genus Chrysochromulina include two species *Chrysochromulina scutellum* and *Chrysochromulin*a sp., both not very clear about their significance in ecosystem. But for the genus Phaeocystis, *P. globosa* was obviously detected among the samples, and this was in accord with our traditional monitoring approaches for the findings of *P.globosa* bloom in Beibu Gulf. Our result convincingly implicated that algae community reflected by eDNA and 18S ribosomal DNA analysis of the V4 region using the Illumina-Based Sequencing platform was suitable for monitoring the harmful algae in the early bloom stage. Many scientist had utilized HTS or NGS to detect algae in aquatic ecosystems and got some attractive results; for instance, NGS had been employed to microalgal diversity in the lichen Ramalina farinacea [42], Diatom resting stages in surface sediments [43], Diatom biomonitoring [40] and detection of harmful algal bloom species [44]. All the results show that eDNA and HTS sequencing is a promising approach to explore the community of algae in aquatic environments [45].

### Hypothesis about the bloom mechanism of *P.globosa* in Beibu Gulf

Some mechanisms about the bloom of *P. globosa* could be proposed from our present work.

The first and most interesting finding was the bloom mode of *P. globosa*. In this study, at the early bloom stage, *P. globosa* was only obviously detected at site S8 with relatively much higher reads (2150) than other sixteen sites (148 in all). Therefore, the bloom of *P. globosa* may originate from a point of site (S8) and then spread to other regions, generally speaking, just as “diffusion from point to face”. Several evidences supported our opinion. Firstly, the marine environment in the Beibu Gulf was protected relatively better than other coastal zones in China [46]. Large area pollution, especially excessive nutrient concentration, had never appeared and been reported previously. Inversely, the probability of point source pollution in the coastal zone was even greater. The first appearance and flourish of *P. globosa* only at site S8 was probably because of its special condition, such as aquaculture farming nearby and consequent eutrophication phenomenon, rich nutrients brought by bottom-up stream and terrestrial drainage. Then the Phaeocystis was carried by transport of some water and drifted along the coastline under the influence of stream in Beibu Gulf. Some previous research could support our opinion. For instance, during the Phaeocystis bloom of the year 1957 around the coast of North Wale, Phaeocystis was only able to proliferate in Liverpool Bay, and then spread to other regions [47]; when Phaeocystis bloom in the coastal of north-western English Channel in 1990, Phaeocystis bloom emanated from near-shore, then spread towards the south-east in accord with the wind direction [48]. If our hypothesis is correct, releasing effective bio-agents and environmental controls on certain region to inhibit harmful algal blooms (HABs) *P. globosa* will be promising [49-51]. In this way, we only need to paid more attention to the early stage of *P. globosa* bloom, not after outbreaks in large scale, and this should be extremely effective.

The second interesting finding about the bloom of *P. globosa* was in relevant to the concentration of NO_3_^-^-N, pH, temperature and DO of seawater. More specifically, the bloom of *P. globosa* has a significant positive correlation with NO_3_^-^-N and negatively related to temperature of seawater. In the Beibu Gulf, the average temperature of seawater was above 15-16 ˚C and the most suitable temperature for growth of *P. globosa* is 15-16 ˚C [52], so the most relative abundance of *P. globosa* appeared first at site S8 may originate from its appropriate temperature of seawater. Nutrient elements, especially nitrogen and phosphorus, have obvious influence on Phaeocystis blooms [52,53]. However, the impact of different forms of N-sources to the bloom of Phaeocystis had never been discovered previously. In our present work, it was the first time that the great correlation between NO_3_^-^-N and Phaeocystis bloom was illustrated. This result provided a helpful reference to our government on how to manage marine environment and control the *P.globosa* bloom. They should pay more attention to reduce the emission of nitrogen, especially NO_3_^-^-N, to the coastal zone of northern Beibu Gulf.

## Acknowledgments

Supported by the National Natural Science Foundation of China (31570875); Guangxi Natural Science Foundation (2014GXNSFBA118135); Foundation of Guangxi Key Laboratory of Marine Disaster in the Beibu Gulf, Qinzhou University (No.2017TS03); Foundation of the Key Laboratory of Coastal Science and Engineering, Beibu Gulf, Guangxi (No.2016ZYB07).

## Author Contributions

### Conceptualization

Bin Gong

### Funding acquisition

Bin Gong, Haiping Wu

### Formal analysis

Bin Gong

### Project administration

Bin Gong, Haiping Wu

### Supervision

Haiping Wu

### Validation

Jixian Ma,Meimiao Luo,Xin Li

### Writing-original draft

Bin Gong

### Writing-review & editing

Haiping Wu

